# Alpha-T-catenin is expressed in peripheral nerves as a constituent of Schwann cell adherens junctions

**DOI:** 10.1101/2022.09.15.508190

**Authors:** Anthea Weng, Erik E. Rabin, Annette S. Flozak, Sergio E. Chiarella, Raul Piseaux Aillon, Cara J. Gottardi

## Abstract

The adherens junction component, alpha-T-catenin (αTcat) is an established contributor to cardiomyocyte junction structure and function, but recent genomic studies link *CTNNA3 polymorphisms* to diseases with no clear cardiac underpinning, including asthma, autism and multiple sclerosis, suggesting causal contributions from a different cell-type. We show *Ctnna3* mRNA is highly expressed in peripheral nerves (e.g., vagus and sciatic), where αTcat protein enriches at paranodes and myelin incisure adherens junctions of Schwann cells. We validate αTcat immunodetection specificity using a new *Ctnna3*-knockout fluorescence reporter mouse line yet find no obvious Schwann cell loss-of-function morphology at the light microscopic level. *CTNNA3/Ctnna3* mRNA is also abundantly detected in oligodendrocytes of the central nervous system via public databases, supporting a general role for αTcat in these unique cell-cell junctions. These data suggest that the wide range of diseases linked to *CTNNA3* may be through its role in maintaining neuroglial functions of central and peripheral nervous systems.

## INTRODUCTION

Alpha-catenins are a family of adherens junction proteins that organize individual cells into tissues through an ability to tether the cadherin/β-catenin cell-cell adhesive complex to actin filaments. They are encoded by separate genes and historically named according to tissue-types where first identified: αE^pithelial^-catenin/*CTNNA1*, αN^eural^-catenin/*CTNNA2*, αT^estes^-catenin/*CTNNA3* (1). αEcat (*CTNNA1*) is now appreciated as broadly expressed and essential for the development and homeostasis of most tissue types (2), leading to it being the most studied form of α-catenin (3–8). αNcat (*CTNNA2*) expression is largely restricted to brain, where it plays critical roles in neuronal synapses required for full brain development (9,10). αT-cat (*CTNNA3*) is the most recently evolved α-catenin, but in contrast to its ancestral relatives (αE-cat and αN-cat) appears developmentally dispensable, as *Ctnna3* knock-out mice are viable and fertile (11,12). Despite its name, αTcat is best known for its role in the heart (12), where it contributes to the interspersed alignment of adherens junctions and desmosomes at the intercalated disk of cardiomyocytes (13–16). This organization is critical for long-term cardiac function, since *Ctnna3* knock-out mice develop age-related cardiomyopathy, similar to patients with loss-of-function mutations in αTcat (13,17). These data have led to the notion that αT-cat plays an important but highly restrictive role in cardiomyocyte functionality.

Curiously, recent genetic association studies have linked mutations, non-coding polymorphisms and copy number variants (CNV) in *CTNNA3* to a wide spectrum of diseases, including asthma, (18–21), food allergy (22), autism spectrum disorder (23–25), multiple sclerosis (26), diabetes (27) and Alzheimer’s (28,29). These disease associations are challenging to rationalize in context of the supposed restricted expression of αT-cat to cardiomyocytes and testes, suggesting αT-cat /CTNNA3 disease linkages may occur via an unappreciated cell/tissue-type. Moreover, while some of the aforementioned genetic studies show genotype/mRNA abundance phenotype correlations (18,20), far fewer show causal changes at the αT-cat protein level, where cross-reactivity of commercially available antibodies between α-catenin family members has been problematic.

Our group previously validated αT-cat’s linkage to steroid-resistant asthma using the house-dust mite model, where *Ctnna3* knock-out mice showed greatly attenuated airway hyperreactivity (30). Despite our observing cardiomyocytes of pulmonary veins as the major αT-cat-expressing cell in lung (31,32), the cell type through which αT-cat loss leads to reduced airway hyperreactivity has remained elusive. Here, we show that αT-cat protein is abundantly expressed in peripheral nerves (e.g., vagus and sciatic), specifically within the Schwann cell component. Since nerves innervate most organ systems and tissue types, we reason that many αT-cat */CTNNA3* disease linkages should be considered via its role in neuroglial cell types.

## RESULTS

Early work on αT-cat/*CTNNA3* used a human cDNA Rapid-Scan™ panel to identify heart and testis as major tissue-types expressing *CTNNA3*, although low levels of RNA were also detected in brain, kidney and liver (2). To better assess the full range of cell- and tissue-types expressing αT-cat */CTNNA3*, we interrogated the Genotype-Tissue Expression (GTEx)-database, an NIH Common Fund resource to study relationships between genetic variation and gene expression across multiple reference tissues (33). This database also serves as a convenient resource to validate RNA abundance across a greater range of tissues (~50-types), especially those not typically harvested in earlier studies (e.g., nerve, sub-regions of brain). This tissue-wide bulk-RNA sequencing database reveals *CTNNA3* expression in tibial nerve, spinal cord and various brain regions (substantia nigra, hippocampus, amygdala) at levels comparable to those observed in heart and testes (https://gtexportal.org/home/gene/CTNNA3). Additionally, single-cell expression within this database reveals *CTNNA3* enrichment in the Schwann cell component of multiple tissues including skeletal muscle, esophagus, prostate and heart. Thus, the GTEx resource reveals that in addition to established expression of in myocytes, *CTNNA3* is also abundantly expressed in the glial component of peripheral nerve.

Since spinal cord and brain register the highest *CTNNA3* expression levels across human tissues, we sought to determine whether this expression might be through a related glial cell-type. To interrogate *CTNNA3* expression in both central and peripheral nervous system, we examined more sensitive RNA-sequencing datasets from enriched (i.e., flow-sorted) cell populations. The Barres Lab RNA-sequencing dataset analyzed gene expression in the cells from the central nervous system, revealing abundant *CTNNA3/Ctnna3* expression in oligodendrocytes and their precursor cells in both humans and mice (34,35). Additionally, the Sciatic Nerve Atlas (SNAT) examined peripheral nerve cell types within mouse sciatic nerve, showing prominent *Ctnna3* expression in Schwann cells (36). Since oligodendrocytes and Schwann cells use cytoplasmic myelin to sheath and thereby insulate neurons of both central and peripheral nervous systems respectively, these databases suggest that αT-cat may support a common role in these functionally related cell types.

### Generation and characterization of a novel Ctnna3-fluorescent reporter mouse

In an attempt to visualize αT-cat expression in cell- and tissue-types not optimally captured by typical paraffin-embedded thin-section methods, we generated a fluorescent reporter mouse in collaboration with Northwestern University’s Transgenic Mouse Core. Using CRISPR-Cas9 gene editing, a membrane anchored enhanced green fluorescent protein (eGFPcaax) (37) was knocked into exon 2 of the *Ctnna3* gene. The *Ctnna3* promoter drives GFP expression while simultaneously knocking out native expression of αT-cat/*Ctnna3* (Fig. 1A). We analyzed eGFPcaax-reporter expression in tissues known to express αTcat protein: heart and testes. We confirmed GFP protein expression in heart tissue from heterozygous (*Ctnna3*^+/eGFPcaax^), but not wild-type (*Ctnna3*^WT^) mice by immunoblot analysis (Fig. 1B). As expected, *Ctnna3*^+/eGFPcaax^ and *Ctnna3*^KO(eGFPcaax/eGFPcaax)^ mice (hereafter referred to as *Ctnna3*^KO^ mice) showed allele-dependent loss of αT-cat protein with complementary changes in GFP abundance in both heart and testes (Fig. S1). While GFP expression from this *Ctnna3*^+/eGFPcaax^ reporter was sufficiently abundant to detect by immunoblotting (Fig. 1B), we were unable to detect GFP by immunofluorescence analysis with an anti-GFP antibody previously validated by our group for use on paraffin-embedded samples, even in *Ctnna3*^KO^ with two copies of GFP (Fig. 1C; GFP staining not shown). Although our GFP reporter contains a membrane-targeting lipid modifiable -CAAX motif to improve detection (37), we reason that localization information embedded within the αT-cat sequence (i.e., ability to bind both β-cat and F-actin and enrich at junction structures) improves its signal-to-noise detection, which the eGFP-caax would lack. Nonetheless, these data show we have generated a GFP-reporter mouse for *Ctnna3* expression (*Ctnna3*^+/eGFPcaax^) that also effectively deletes the endogenous αT-cat protein when crossed to homozygosity (*Ctnna3*^KO^).

**Figure 1:**
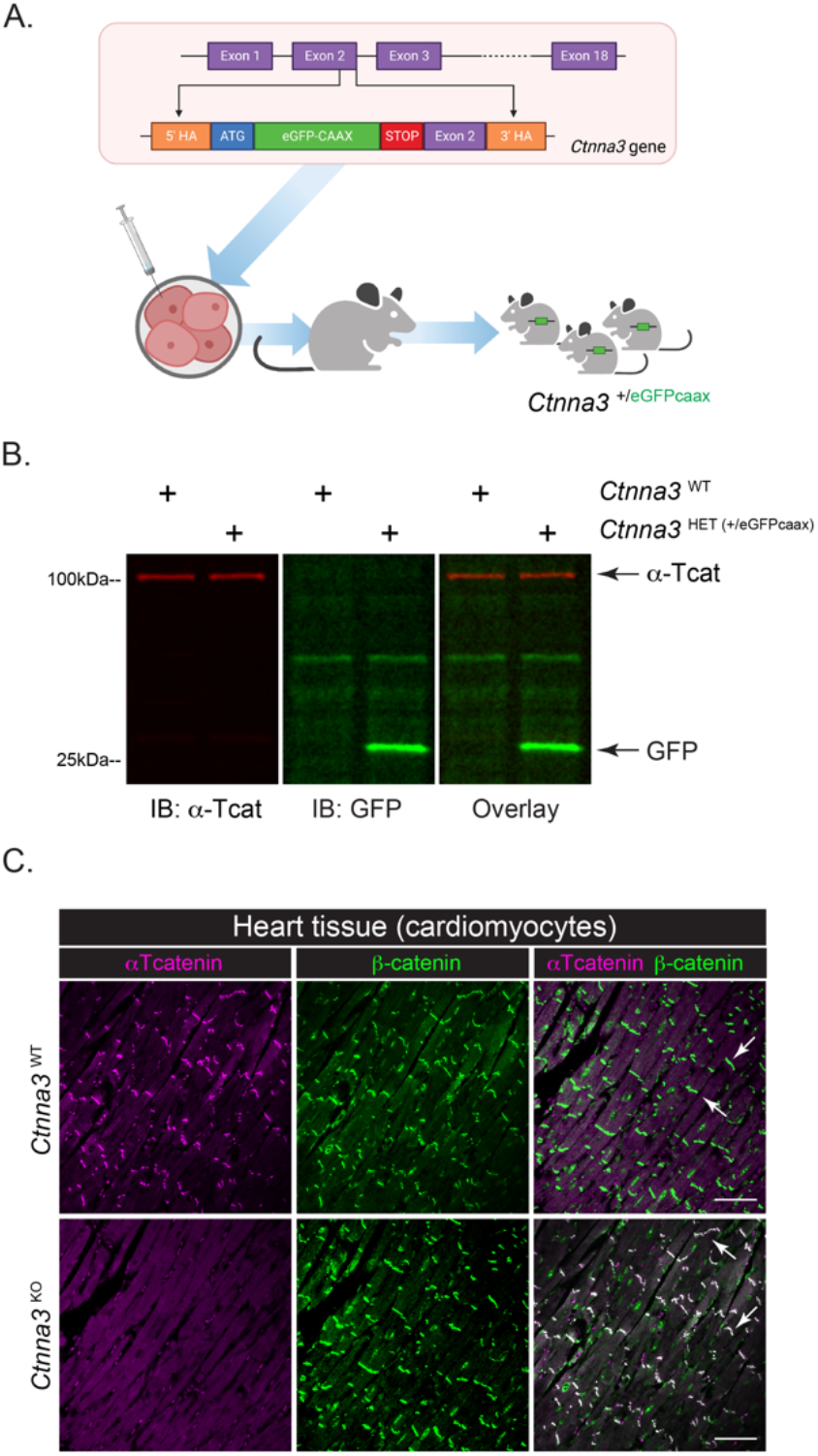
Generation and characterization of a novel fluorescent *Ctnna3-*reporter mouse. (**A**): Reporter design schematic by BioRender. Membrane-anchored eGFPcaax was inserted into exon 2 of the *Ctnna3* gene. Construct was nucleofected into embryonic stem cells, clones with correct construct insertion were selected, and two chimeric males were generated. (**B**): Immunoblotting of *Ctnna3*^WT^ and *Ctnna3*^HET (+/eGFPcaax)^ heart lysates with anti-αTcat (red) and anti-GFP (green) antibodies. (**C**): Representative images of heart sections from *Ctnna3*^WT^ and *Ctnna3*^KO^ mice. Scale bars, 50μm. Immunofluorescence staining of cardiomyocytes by β-catenin (β-cat; green) and αTcat (magenta). Arrows indicate intercalated disk-type adherens junctions in cardiomyocytes.

### αTcat/Ctnna3 is expressed in vagus nerve

A number of genome-wide association studies have linked *CTNNA3* to chemical-induced occupational asthma (18,19), steroid refractory asthma (21) and asthmatic exacerbations (20). Indeed, our team validated αT-cat’s linkage to asthma using the house-dust mite model, where *Ctnna3* knock-out mice showed greatly attenuated airway hyperreactivity (30). However, despite our observing cardiomyocytes of pulmonary veins as the major αT-cat-expressing cell in lung (30–32), the cell type through which αT-cat loss leads to reduced airway hyperreactivity has remained elusive. Given evidence via the GTEx database that *CTNNA3* is abundantly expressed in peripheral nerves, where parasympathetic peripheral nerve inputs are known to innervate airways and control smooth muscle contractility/reactivity responses (38–40) we sought to determine whether αTcat protein might be expressed in the vagus. We harvested small (~0.5cm) segments of the vagus nerve and nodose ganglion just anterior to the thoracic cavity of wild-type (*Ctnna3*^WT^) and knock-out (*Ctnna3*^KO^) mice for total protein and RNA extraction (Fig. 2A, schematic). We confirmed αTcat expression in both nodose ganglion and vagus nerve in wild-type mice; Knock-out tissue showed no αTcat (Fig. 2B). Remarkably, we detected *Ctnna3* RNA expression in vagus and nodose at levels comparable to and even greater than *Ctnna3* levels in heart (Fig. 2C). Moreover, since αTcat/*Ctnna3* RNA is as abundant in vagus segments as in nodose ganglion, where the latter contains neuronal cell bodies, we reason that the bulk of *Ctnna3* signal may be due to its expression in Schwann cells, consistent with database resources showing little *Ctnna3* RNA in neurons (34,35).

**Figure 2:**
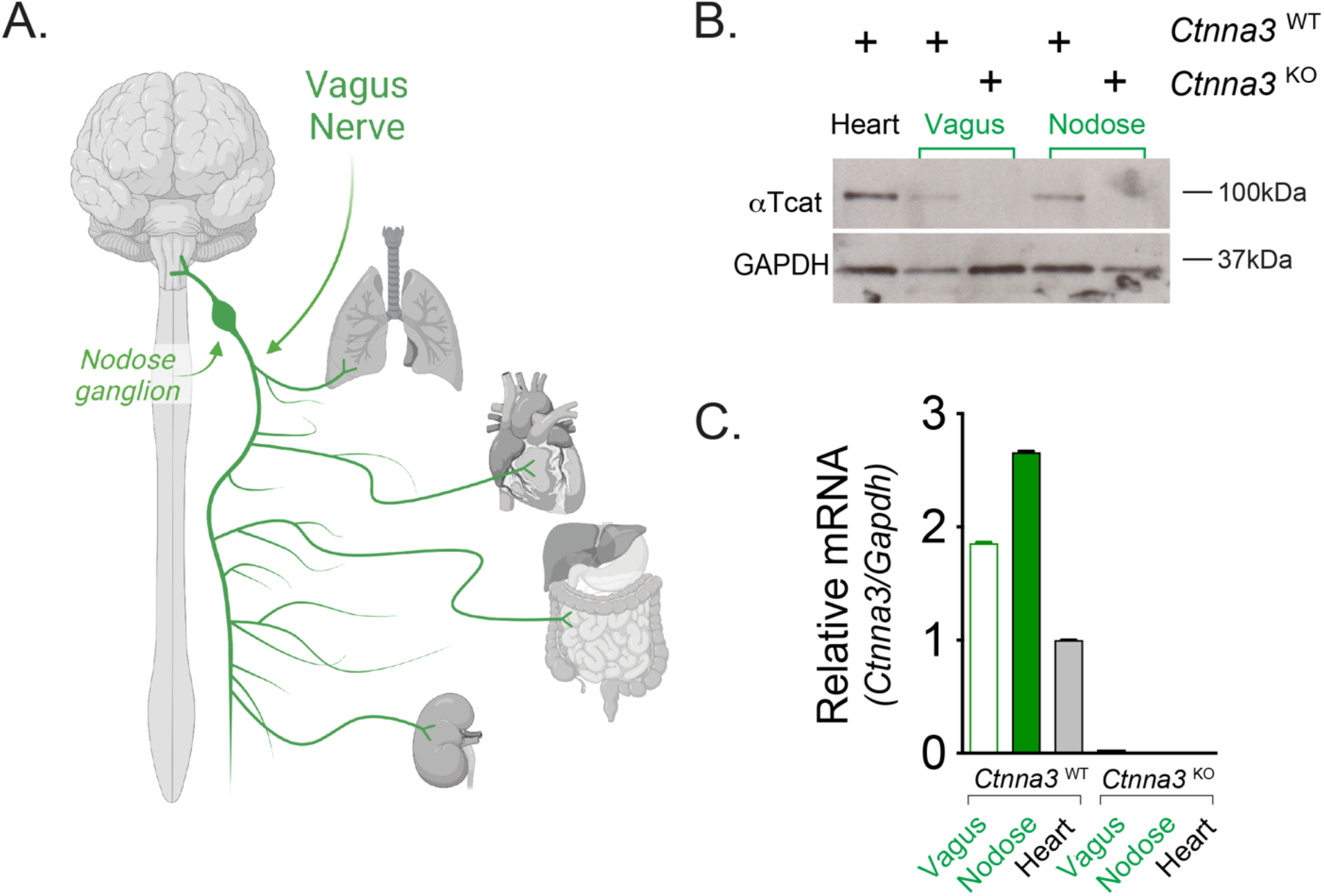
αTcat is expressed in vagus nerve. (**A**): Schematic of nodose ganglion, vagus nerve and organs innervated by vagus (Biorender). (**B**): Western blot of *Ctnna3*^WT^ and *Ctnna3^KO^* heart, vagus and nodose tissue with anti-αTcat and loading control GAPDH. (**C**): mRNA relative expression via real-time quantitative PCR analysis of *Ctnna3*^WT^ and *Ctnna3^KO^* RNA isolated from heart, vagus and nodose tissues. Error bars reflect technical (n = 3) rather than biological replicates due to difficulty harvesting tissue and independent validation of αTcat protein in B.

### αTcat is expressed along Schwann cell myelin incisures of sciatic nerve

To specifically address whether αTcat protein is expressed in the Schwann cell component of peripheral nerves, we turned to the sciatic nerve due to its large size and ease of dissection (Fig. 3A, Supplemental Video 1). We validated αTcat protein expression in sciatic nerve lysates harvested from *Ctnna3*^WT^ but not *Ctnna3*^KO^ mice; the ubiquitous αEcat protein remained largely unchanged (Fig. 3B). Immunofluorescence staining of wild-type sciatic nerve revealed αTcat localization to paranodal regions of glial-axon junctions, as well as structures reminiscent of myelin incisures, thin regions of Schwann cell cytoplasm excluded from compact myelin regions (Fig. 3C). Also known as Schmidt-Lanterman incisures, this structure results from successive concentric wraps of Schwann cell cytoplasm around an axon, where each wrapping is held together by adherens junction proteins in a uniquely characteristic intracellular “autotypic” junction (41,42). As expected, αTcat protein co-localizes with known adherens junction markers, E-cadherin and F-actin (Fig. S2). Importantly, *Ctnna3*^KO^ mice showed no detectable αTcat immunostaining (Fig. 3C, bottom and 3D, right). We observed no obvious structural defect in these nerves, as overall E-cadherin and F-actin staining patterns were similar. Disappointingly, we were also unable to rely on GFP immunostaining as a convenient reporter for *Ctnna3* expression in this tissue, even in *Ctnna3*^KO^ mice with two copies of eGFPcaax (not shown). In summary, these data show that αTcat protein localizes to Schwann cell autotypic and heterotypic junction structures with familiar partners, E-cadherin and F-actin.

**Figure 3:**
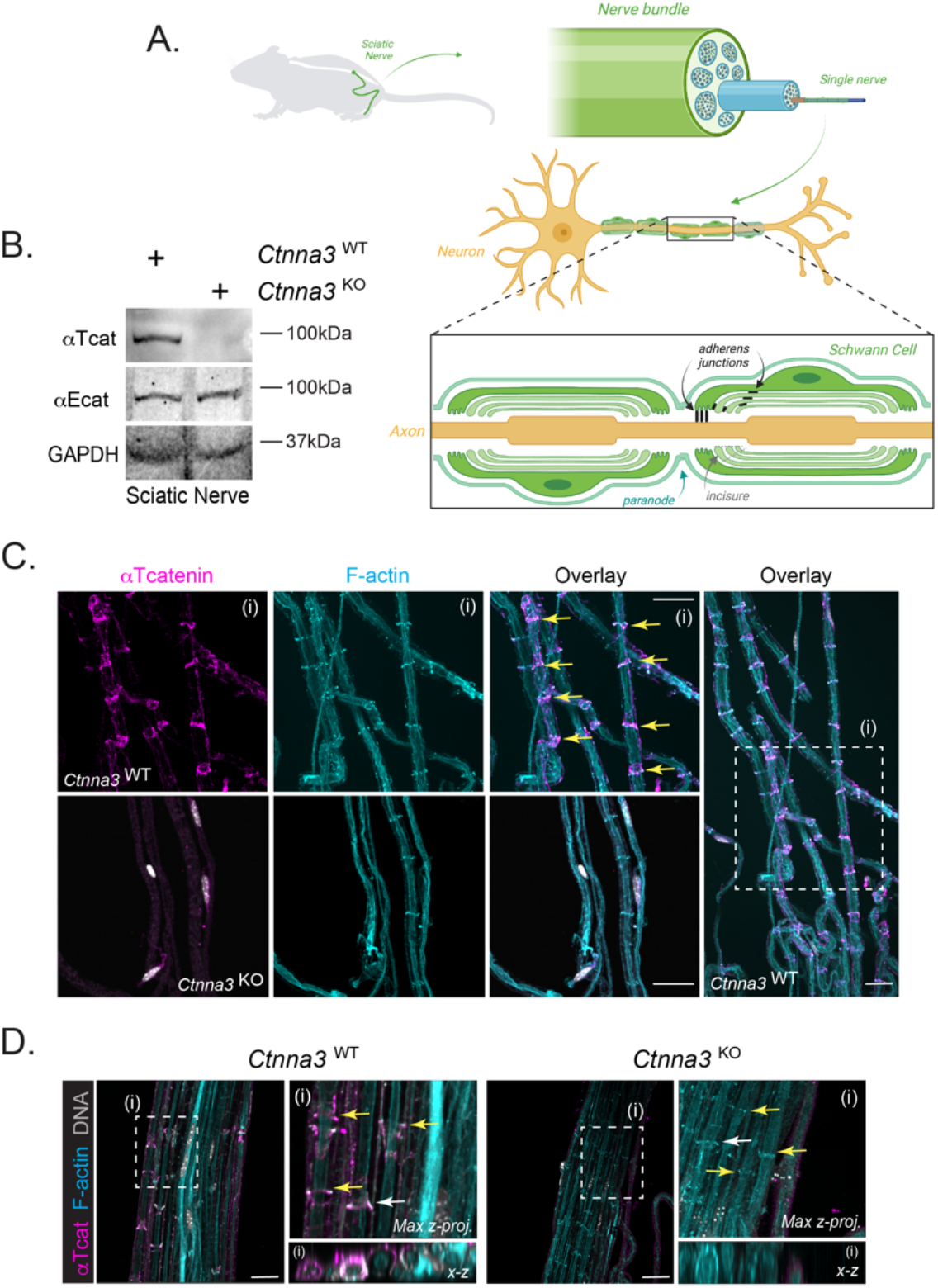
αTcat is expressed at myelin incisure and paranodal loop regions of Schwann cells in sciatic nerve. (**A**): Schematic (Biorender) demonstrating αTcat expression at adherens junctions along myelin incisures and paranodal loops of Schwann cells from sciatic nerve. (**B)**: Western blot of *Ctnna3*^WT^ and *Ctnna3^KO^* sciatic nerve tissue with anti-αTcat, anti-αEcat, and loading control GAPDH. (**C-D**): Representative confocal images of sciatic nerve sections from adult (8-12 weeks) from *Ctnna3*^WT^ and *Ctnna3*^KO^ mice; one field of view from low- and high-magnification inset (boxed region) are shown (box i). Low-magnification view for Ctnna3^KO^ is not shown. Immunofluorescence staining of myelin incisures by αTcat (magenta) and F-actin (cyan). DNA labeling by Hoechst (gray). Scale bars, 25μm. Yellow arrows indicate adherens junctions where αTcat co-localizes with F-actin. (D): Maximum intensity z-projection images were generated with Imaris from confocal images. White arrows indicate plane of x-z section.

## DISCUSSION

αTcat is the most recently evolved member of the alpha catenin family and best known for its role in cardiomyocyte junctions of heart, where its loss leads to an age-related cardiomyopathy that phenocopies disease in patients with αTcat loss-of-function mutations (12). These data have led to the notion that αT-cat plays an important but highly restrictive role in cardiomyocyte functionality. However, numerous genetic association studies have linked *CTNNA3* to a wide spectrum of diseases, ranging from asthma and food allergy to autism, multiple sclerosis, diabetes and Alzheimer’s (18–29). These disease associations have been puzzling to rationalize in context of established restricted expression of αT-cat to cardiomyocytes (or testes), suggesting αT-cat/*CTNNA3* disease linkages may occur via an unappreciated cell/tissue-type. Guided by the human tissue gene expression (GTEx) database, we demonstrate αTcat protein is also highly expressed in peripheral nerves (e.g., vagus and sciatic), specifically within the Schwann cell component. *CTNNA3/Ctnna3* mRNA is also abundantly expressed in oligodendrocytes of the central nervous system (34,35). Together with previous evidence that αTcat localizes to apical junctions of ependymal cells (43), we propose a generalized role for αT-cat in neuroglial functions. Since neuroglia participate in central and peripheral nervous systems, where the latter innervate all organs of the body, we speculate that the wide range of αT-cat/*CTNNA3* disease linkages may be considered via its role in neuroglial cell types.

Anticipating the need to visualize *Ctnna3* gene expression in a cell type challenging to resolve by thin section imaging (e.g., Schwann cells), we generated a novel fluorescent reporter *Ctnna3*^+/eGFPcaax^ knockout mouse. While we validated this reporter line for GFP expression and consequent loss of αTcat, we were unable to visualize the membrane-anchored GFP reporter by immunofluorescence analysis. Whether this is due to sequences in *Ctnna3* that enhance the translation efficiency of αTcat, or the ability of αTcat to become highly enriched at cell-cell junctions compared with a general membrane-localized GFP is not clear. Nonetheless, this reporter may ultimately prove valuable for future studies seeking to characterize consequences of *Ctnna3*/αTcat loss in neuroglial cell types.

It is intriguing that αTcat/*CTNNA3* expression is restricted to Schwann cells of peripheral nerves and oligodendrocytes in brain, two cell-types known to play critical roles in the myelination and insulation of neurons. Our evidence that αTcat specifically localizes to paranodes and myelin incisures of Schwann cells was not entirely surprising, given previous evidence that both structures comprise modified adherens junctions enriched for cadherins and catenin proteins (41,42). Paranodes reflect regions of heterotypic glial cell-axon contact, whereas Schmidt-Lanterman myelin incisures reflect an alignment of successive wraps of Schwann cell cytoplasm around an axon, where each wrap is held together by adherens junction proteins (41). While forced-expression of dominant-inhibitory versions of cadherin-catenin complex components can perturb this structure (42), whether polymorphisms or mutation in members of this cadherin-catenin adhesive complex can alter the critical function of Schwann cells (and related oligodendrocytes) relevant to disease remains unknown.

Our evidence that αTcat protein is detected in the vagus, which innervates the majority of visceral tissues throughout the body, offers an attractive lens through which we may view genetic linkages between αTcat/*CTNNA3* and a range of diseases (reviewed in (1)). With regards to *CTNNA3* linkages to steroid-resistant asthma, our team previously validated this association showing that *Ctnna3*^KO^ mice display reduced airway hyperreactivity in response to methacholine challenge. Our evidence that αTcat is expressed in the vagus, a known regulator of airway smooth muscle responses (38–40), suggests the intriguing possibility that αTcat/*CTNNA3* contributes to asthmatic airway responses through its role in the Schwann cell-myelinating component of peripheral nerves. Indeed, even *CTNNA3*’s linkages to heart disease merit revisiting, where recent linkages to atrial fibrillation/Brugada Syndrome (44) may be due to a peripheral nerve rather than a cardiomyocyte junction defect (45).

## METHODS

### *Ctnna3* membrane-anchored GFP reporter mouse

We knocked-in a membrane associated eGFP construct into exon2 of the *Ctnna3* locus using CRISPR gene editing in collaboration with Northwestern’s Transgenic and Targeted Mutagenesis Laboratory. By design, the *Ctnna3* promoter drives eGFP-caax expression from the endogenous locus, while knocking out expression of the gene. Note, addition of an exogenous bGH polyA immediately downstream of the stop codon was required to improve eGFP-caax expression from the endogenous *Ctnna3* exon 2 locus, given the extremely large size of the Ctnna3 gene. We nucleofected the following into murine B6N-derived embryonic stem (ES) cells: *Ctnna3*-targeting CRISPR reagents, an eGFP repair plasmid designed to insert into the *Ctnna3* exon2 locus, and a non-targeting PGK-plasmid harboring a puromycin selection cassette. Puromycin resistant ES cell clones were selected, propagated and genotyped for correct construct insertion. Targeted clones were expanded and validated, with select clones injected into albino B6J blastocyst stage embryos. Briefly, donor females (albino B6J) are hormone treated, mated, and plugged. Recipient (foster) females are also set up. ES cells are injected into the cavity of an expanded blastocyst stage embryo and injected embryos are surgically transferred into the reproductive tract of recipient females to generate chimeric mice. Chimeric mice were genotyped for correct insertion of the construct into the *Ctnna3* locus. Two chimeric males (90% and 95%) were generated and bred to C57BL/6J mice, transmitting GFP to the F1 generation with initial genotype validation by endpoint PCR and currently via real-time PCR with Transnetyx (Cordova, TN). GFP expression was validated by immunoblot/I mmunostaining analysis of target tissues. Detailed mouse report with PCR-gel validation will be provided upon request. All experimental protocols were approved by the Institutional Animal Care and Use Committee at Northwestern University.

### Tissue collection

Mice were euthanized and perfused with 10 mL of HBSS. For heart dissection, tissue was fixed in 4% PFA overnight at 4°C, then transferred into 70% ethanol for dehydration, paraffin embedding and ~4μm sectioning via microtome. For vagus nerve and nodose ganglion dissection, skin, salivary glands, and masticatory muscles were removed, followed by identification of the trachea, carotid artery, and glossopharyngeal nerve. The vagus nerve and nodose ganglion were identified, excised, and stored at −20°C for further processing as previously described (46). For sciatic nerve dissection, skin and muscle of upper hind legs were removed, and a ~3mm segment of sciatic nerve was cut and placed into Zamboni’s fixative for 10 minutes, followed by 15% glycerol/PBS solution (v/v) for 24 hours at 4°C. Nerves were then incubated in successive glycerol solutions (45%, 60%, 66% glycerol in PBS) for 18-24 hours each at 4°C. Tissue was stored at −20°C until ready for further processing. Nerve bundles were further dissected into individual fibers using a Leica MZFLIII dissecting microscope. Briefly, nerves were placed into 10cm dishes containing HBSS. Nerves were teased apart using 30G needles and individual fibers were placed onto poly-L-lysine-coated Bond380 slides. Slides were then dried and stored at −20°C.

### Immunofluorescent staining and Imaging

For paraffin-embedded heart slides: Slides were de-paraffinized by submerging in xylene twice for 5 minutes each. Slides were then gradually rehydrated in 100%, 90%, and 70% EtOH solutions twice for 2 minutes followed by three washes in PBS for 5 minutes each. Slides were subsequently quenched in a 10 mM glycine solution for 15 minutes at room temperature. Slides were simmered in a citrate-based antigen retrieval buffer for 30 minutes at 95°C. After cooling to 25°C, slides were briefly rinsed in PBS before immunostaining. For heart and sciatic nerve tissue, slides were blocked in a solution of 10% normal goat serum (NGS) in 0.3%TritonX-100 PBS for 30 minutes at 25°C. Slides received primary antibody prepared in a solution containing 3% NGS in 0.3%TritonX-100 PBS and incubated for 1 hour at 25°C. Following primary antibody, slides were rinsed in PBS and then incubated in fluorescence conjugated secondary antibody for 30 minutes at 25°C. Tissue sections were briefly rinsed in water and mounted using ProLong Gold anti-fade mounting media. Heart tissue was imaged on a Zeiss Axioplan2 epifluorescence microscope. Sciatic nerves were imaged at Northwestern’s Center for Advanced Microscopy core using a Nikon W1 spinning disk confocal microscope. 20x z-stack images were taken in 0.5μm steps whereas 60x z-stack images were taken in 0.25μm step sizes. All image files were pseudo-colored in FIJI for presentation in figures. 3D imaging of sciatic nerve was analyzed using Imaris software. Orthogonal views were obtained from maximum intensity projection (MIP) images. Videos were created via 3D reconstruction of z-stacks (10-25μm depth; 0.25μm step size).

### Tissue preparation and Western blotting

Heart, vagus/sciatic nerve and nodose ganglion tissues were collected and snap-frozen in liquid nitrogen. Heart tissue was homogenized on ice using Tissue Tearor at medium speed for 1 minute in lysis buffer (RIPA plus 0.1% SDS with Roche EDTA-free protease inhibitor) and allowed to sit on ice 10 minutes followed by sonication using Branson sonifier at 10% amplitude, 1 second on/off intervals for 10 rounds total. Lysates were centrifuged at 14,000g for 10 minutes at 4°C and the supernatant collected and stored at −80°C. Vagus nerve and nodose tissues were processed as described except that tissues were pipetted up and down 20 times in lysis buffer using a large bore tip and allowed to sit on ice 10 minutes before sonication and centrifugation. Protein concentrations were determined via Bradford assay. Samples were diluted in 2x SDS loading buffer and boiled at 95°C for 5 minutes prior to loading. 50μg of protein was run on a 4-20% gel at 120V for 90 minutes. Following gel transfer, nitrocellulose membranes were rinsed in Ponceau S to confirm presence of protein. Membranes were blocked for 1 hour at room temperature in 5% milk/TBS-T, followed by incubation in primary antibody in 5% BSA/TBS-T overnight with rocking at 4°C. After primary antibody incubation, membranes were washed 3 times for 10 minutes in TBS-T. Membranes received secondary antibody (HRP or fluorescent conjugated) prepared in a 5% milk/TBS-T solution for 1 hour at room temperature with rocking. Membranes were then washed 3 times for 15 minutes in TBS-T before either incubation with Pierce ECL Plus solution for 1 minute or direct fluorescence imaging on Licor Odyssey Fc scanner for 2-10 minutes.

### RNA isolation and PCR analysis

RNA was extracted from snap-frozen heart, vagus nerve, and nodose ganglion tissue using the Qiagen RNeasy Plus Mini Kit as per manufacturer instructions. Briefly, tissue samples were lysed in 600ml RLT buffer using sterile Beadbug homogenization tubes with 3mm ceramic beads and Benchmark Beadblaster shaker run at power setting 4, 30 second shake with 30 second intervals, 4 cycles, program repeated twice. Homogenized sample was centrifuged at 14,000g for 3 minutes. Supernatant was applied to Qiagen gDNA Eliminator spin columns before continuing with Qiagen protocol. RNA was eluted in 40ml nuclease-free water. First strand cDNA was reverse transcribed from equal amounts of total RNA using iScript cDNA synthesis kit (Biorad, Hercules, CA). Specific primers for amplification of the mouse *Ctnna3* message: Forward Primer (FP) 5’-GGTTACTACCCTGGTGAATTGTCC-3’, Reverse Primer (RP) 5’-CTCTTTTCGAACTTCCTGGAGTGC-3’. Real-time PCR was performed using iQ SYBR Green Supermix according to the manufacturer instructions (Biorad, Hercules, CA). PCR was carried out in 96-tube plates using the MyiQ Single Color Real-Time PCR Detection System and software (Bio-Rad). All reactions were done in triplicate with negative controls. Gapdh was used as the internal control. The relative change in gene expression was calculated using the 2^ΔΔCt^ method.

## ACKNOWLEDGEMENTS

We thank Bruce Appel (U. Colorado, Denver) for the membrane-anchored eGFPcaax construct and Lynn Doglio, Eugene Wyatt and Rajesh Awatramani at Northwestern University’s Transgenic Mouse Core for advice and design of the *Ctnna3*^eGFPcaax^ reporter mouse line. We thank Robert P. Schleimer (Allergy and Immunology, Northwestern) and Bradley Undem (Johns Hopkins University) for discussions regarding vagal contributions to asthma. CJG was supported by Northwestern University Allergy Immunology Research Program (NUAIR; T32AI083216), GM129312, Center for Advanced Microscopy (NCI CCSG P30 CA060553; S10 RR031680; S10OD016342).

## Author Contributions

AW, EER, SEC, ASF and RPA designed and conducted experiments; AW, SEC, ASF and CJG analyzed results. RPA and SEC isolated sciatic and vagus nerves, respectively; AW performed immunofluorescence, imaging analysis and mouse colony maintenance; EER carried out PCR analysis of vagus. CJG designed and supervised study, performed analysis and wrote manuscript. CJG provided funding for project.

---

**Figure S1:**
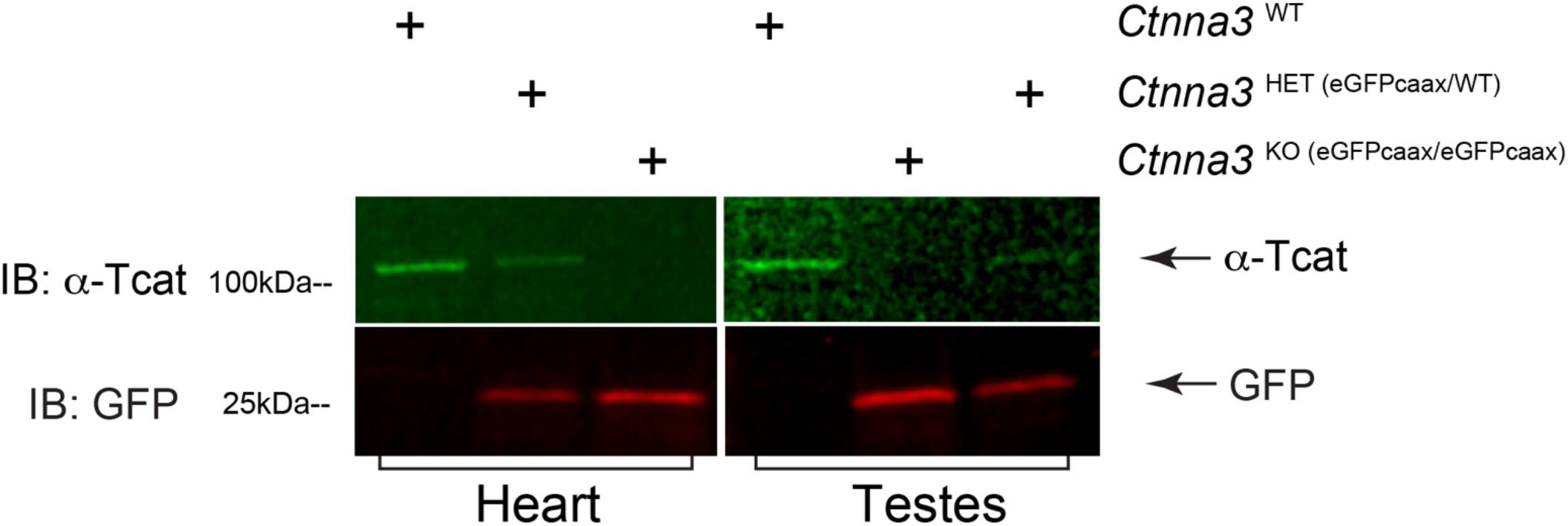
Validation of *Ctnna3*^eGFPcaax^-reporter mouse GFP expression in heart and testes. Immunoblotting of *Ctnna3*^WT^, *Ctnna3*^HET (+/eGFPcaax)^ and *Ctnna3*^KO (eGFPcaax/eGFPcaax)^ lysates from heart (left) and testes (right) with anti-αTcat (green) and anti-GFP (red) antibodies. Note copy-dependent increase in GFP abundance between *Ctnna3*^HET (+/eGFPcaax)^ and *Ctnna3*^KO (eGFPcaax/eGFPcaax)^ mice, with complementary reduction in αTcat protein.

**Figure S2:**
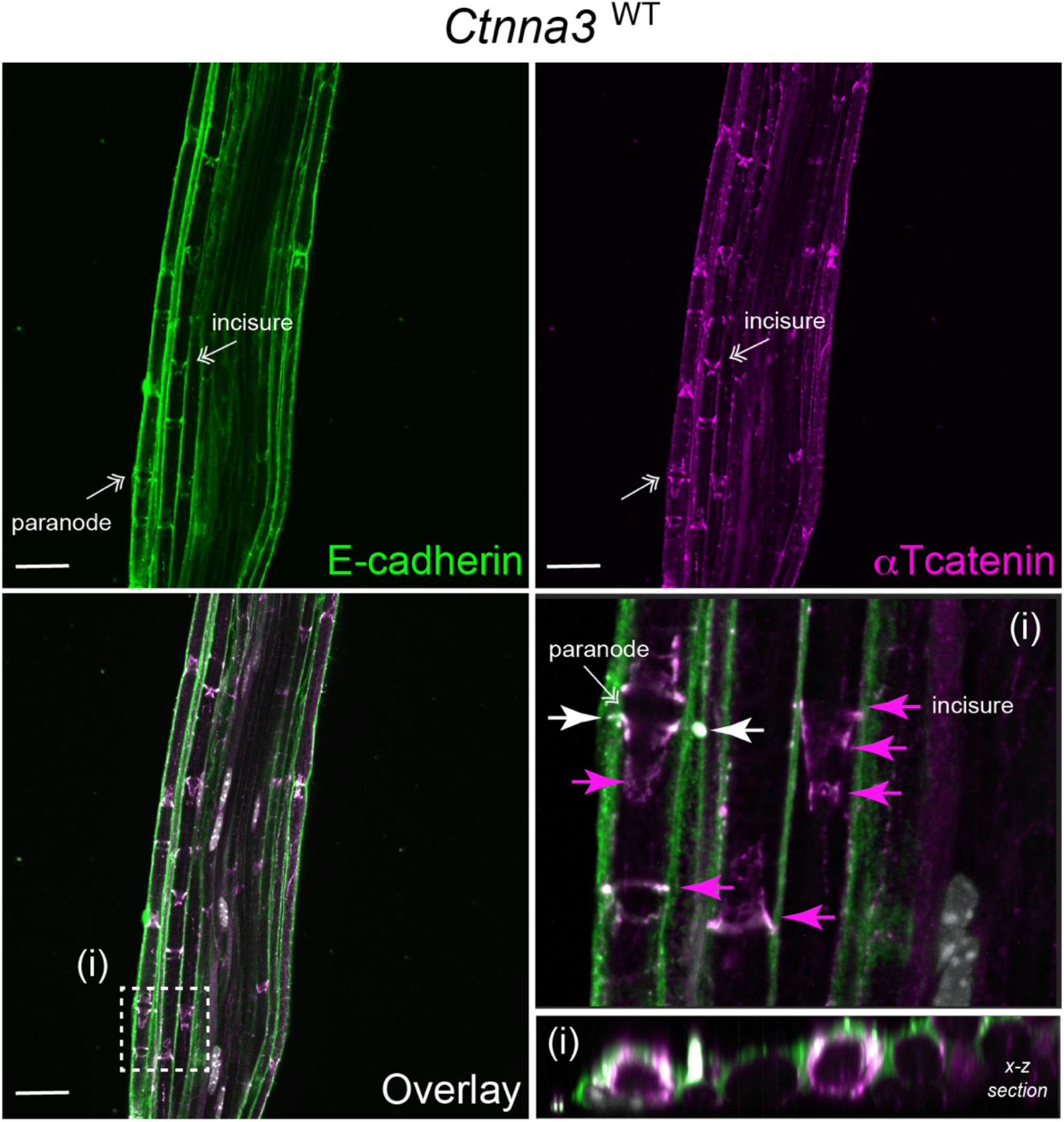
*Ctnna3* co-localizes with known junction marker, E-cadherin. Representative confocal images of sciatic nerve sections from adult (8-12 weeks) wild-type (*Ctnna3*^WT^) reporter mice; one field of view from low- and high-magnification insets (boxed region) are shown (box i). Immunofluorescence staining of adherens junction in myelin incisures by E-cadherin (green) and αTcat (magenta). DNA labeling by Hoechst (gray). Scale bars, 25μm. White arrows indicate plane of x-z section. Areas of E-cadherin and αTcat co-localization are revealed by white fluorescent staining, showing αTcat co-localizes with known junction markers at myelin incisures in sciatic nerve.

**Video 1: αTcat is expressed at myelin incisures in sciatic nerve.** Low- and high-magnification Z-stacks (0.25μm steps; 10μm depth) of sciatic nerve sections from *Ctnna3*^WT^ and *Ctnna3*^KO^ were reconstructed in 3D using Imaris. Immunofluorescence staining of myelin incisures by αTcat (magenta) and F-actin (cyan). DNA labeling by Hoechst (gray).

## KEY RESOURCES

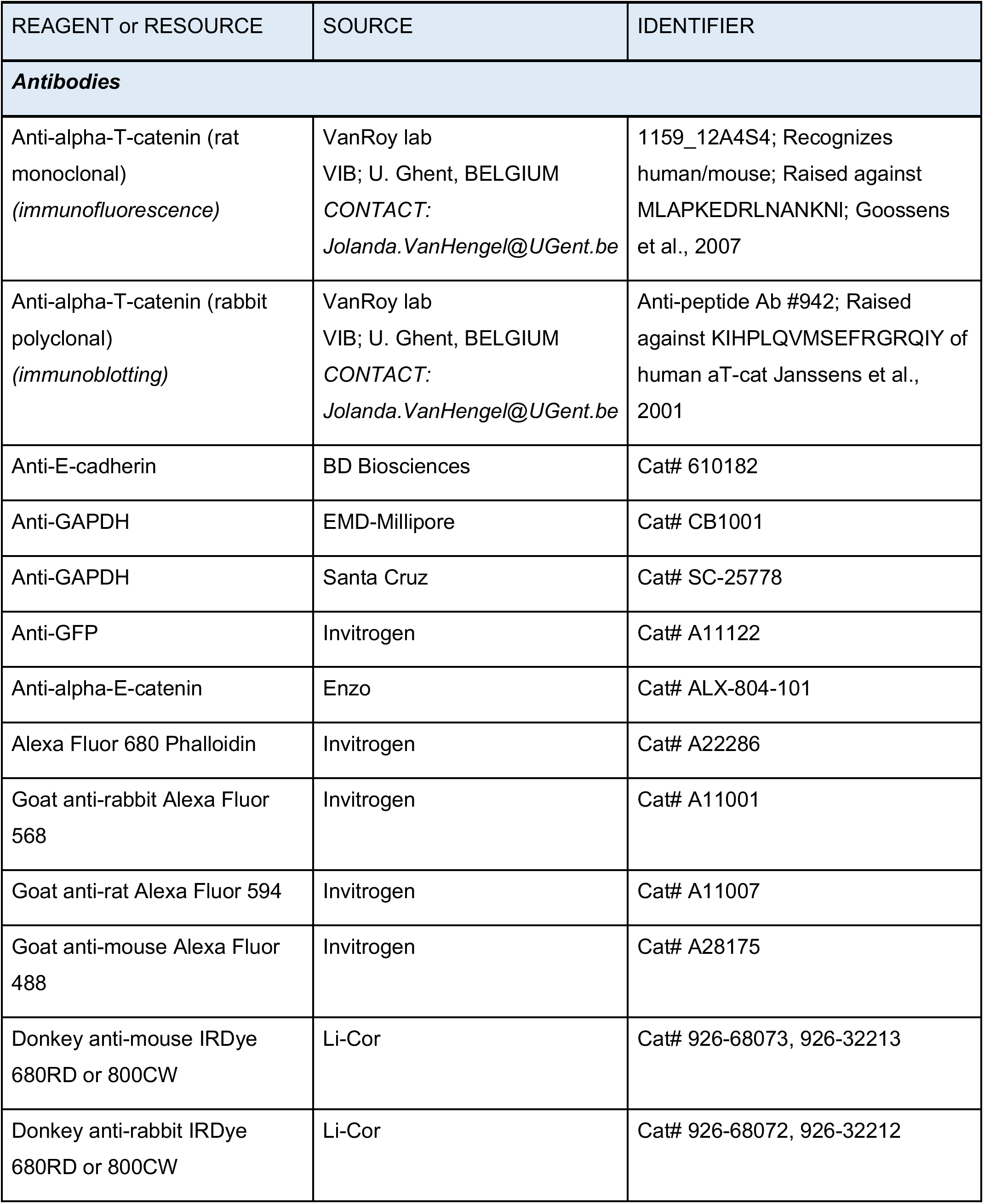
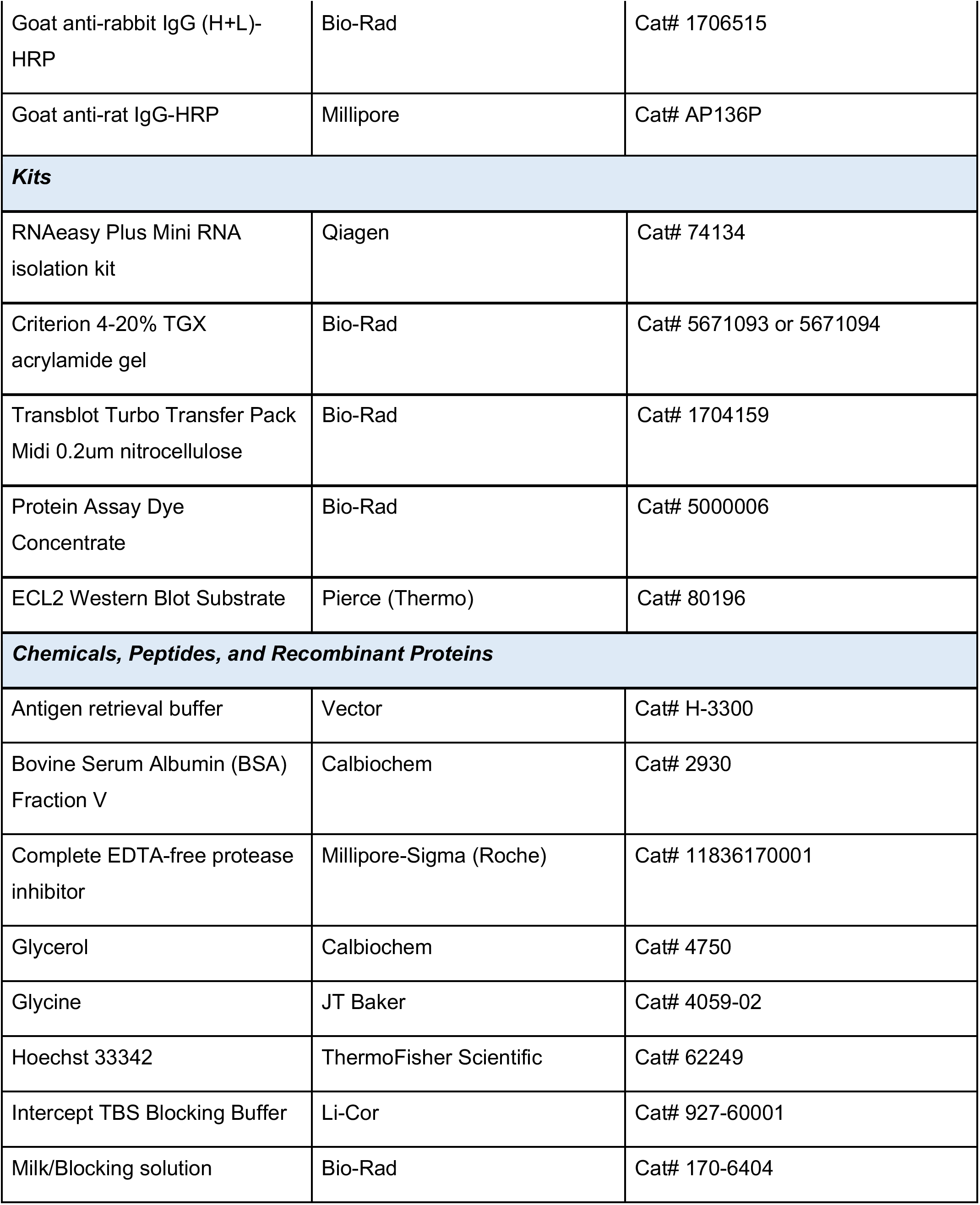
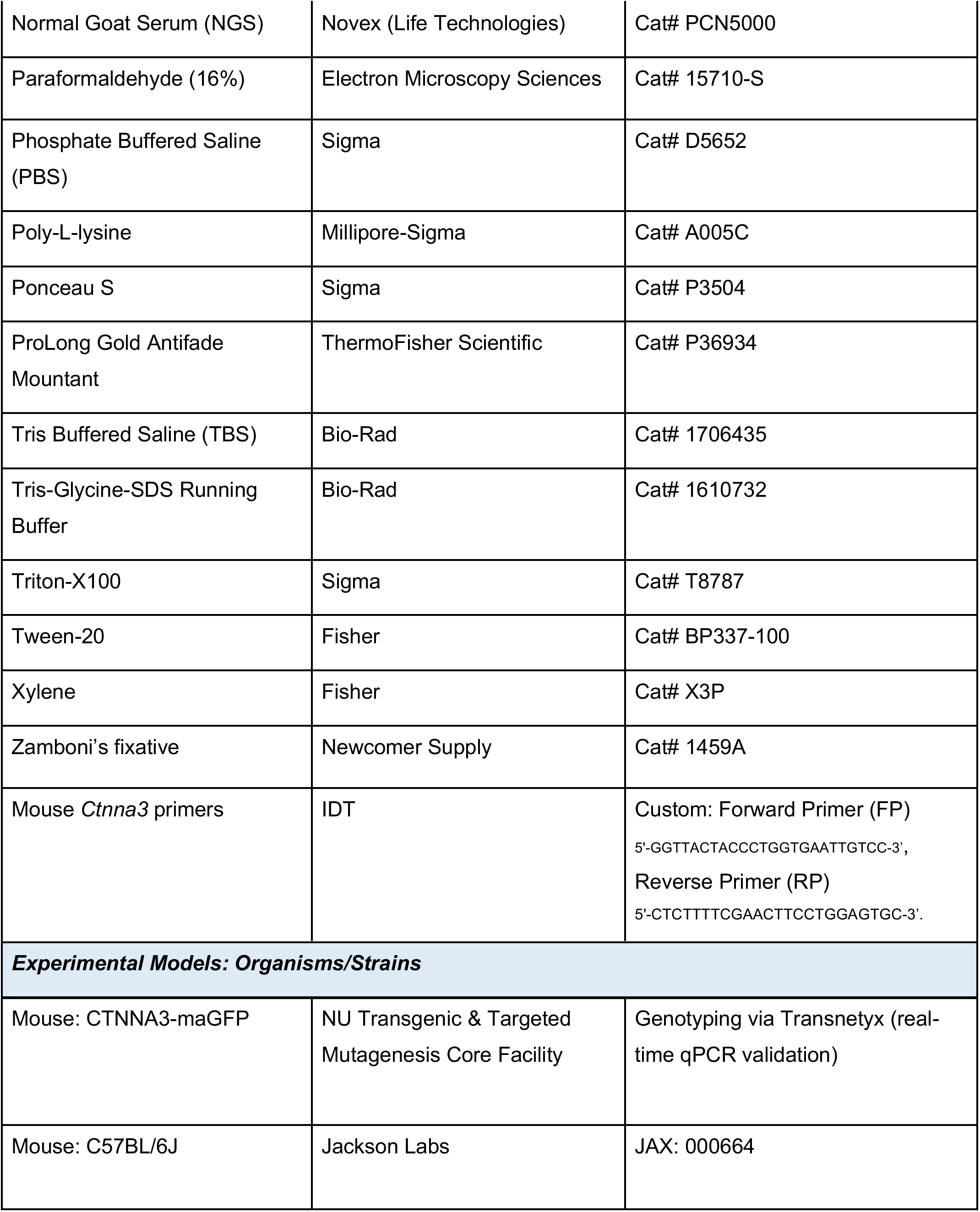
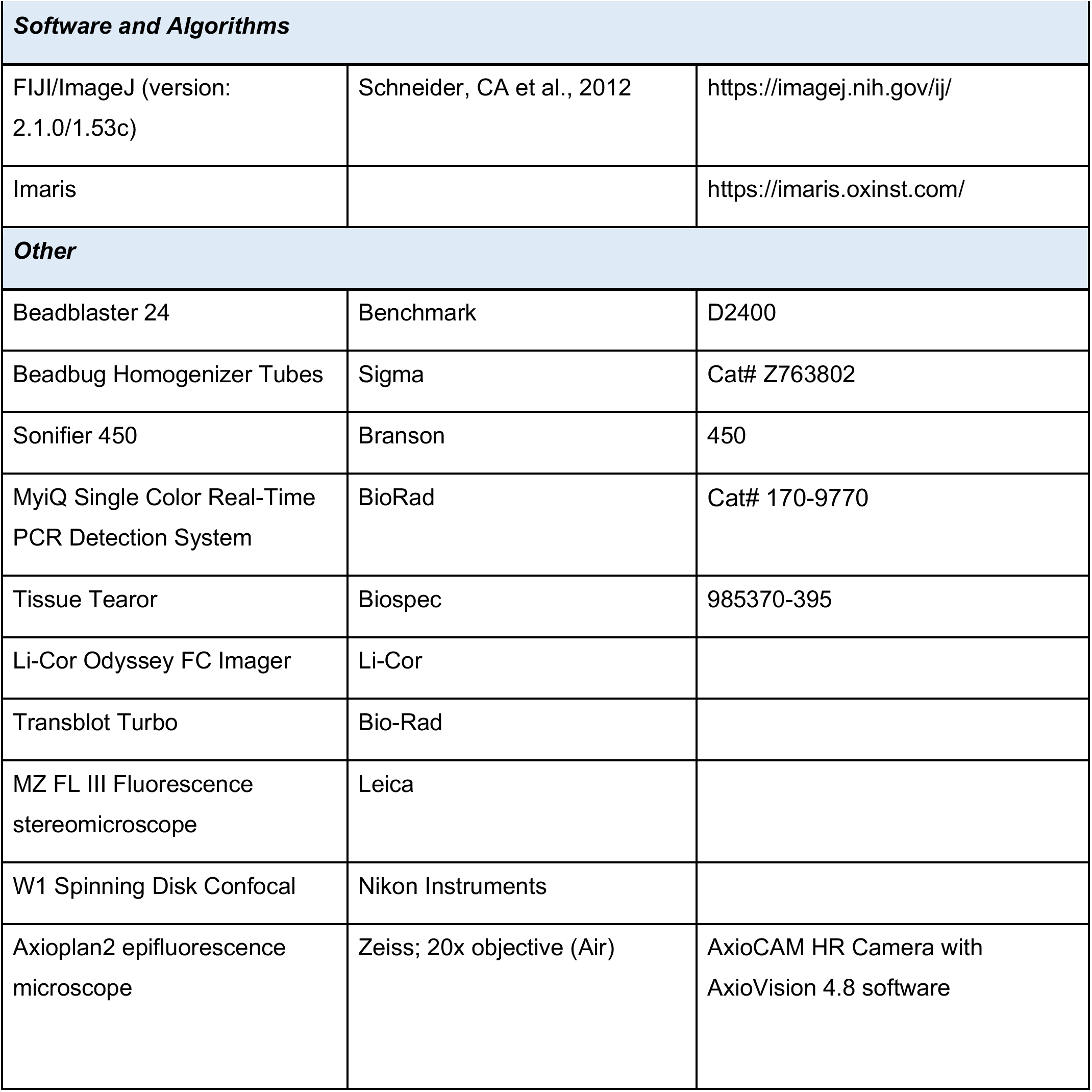

## Notes

### Competing Interest Statement

The authors have declared no competing interest.

